# Molecular coevolution of nuclear and nucleolar localization signals inside basic domain of HIV-1 Tat

**DOI:** 10.1101/2021.04.20.440437

**Authors:** Margarita A. Kurnaeva, Arthur O. Zalevsky, Eugene A. Arifulin, Olga M. Lisitsyna, Anna V. Tvorogova, Maria Y. Shubina, Gleb P. Bourenkov, Maria A. Tikhomirova, Daria M. Potashnikova, Anastasia I. Kachalova, Yana R. Musinova, Andrey V. Golovin, Yegor S. Vassetzky, Eugene V. Sheval

## Abstract

During evolution, viruses had to adapt to an increasingly complex environment of eukaryotic cells. Viral proteins that need to enter the cell nucleus or associate with nucleoli possess nuclear localization signals (NLSs) and nucleolar localization signals (NoLSs) for nuclear and nucleolar accumulation, respectively. As viral proteins are relatively small, acquisition of novel sequences seems to be a more complicated task for viruses than for eukaryotes. Here, we carried out a comprehensive analysis of the basic domain (BD) of HIV-1 Tat to show how viral proteins might evolve with NLSs and NoLSs without an increase in protein size. The HIV-1 Tat BD is involved in several functions, the most important being the transactivation of viral transcription. The BD also functions as an NLS, although it is substantially longer than a typical NLS. It seems that different regions in the BD could function as NLSs due to its enrichment with positively charged amino acids. Additionally, the high positive net charge inevitably causes the BD to function as an NoLS through a charge-specific mechanism. The integration of NLSs and NoLSs into functional domains enriched with positively charged amino acids might be a mechanism that allows the condensation of different functional sequences in small protein regions and, as a result, to reduce protein size, influencing the origin and evolution of NLSs and NoLSs in viruses.

---

The origin of the cell nucleus, which led to the emergence of eukaryotic cells, was one of the most fateful events during the evolution of life on Earth. Acquisition of a nucleus enabled the spatial segregation of transcription and translation and led to the evolution of sophisticated mechanisms of regulation of gene expression (1).

The origin of the nuclear envelope led to the evolution of a complex system for nuclear import of proteins that are produced in the cytoplasm. To enter the nucleus, proteins use short signal sequences, known as nuclear localization signals (NLSs) (2, 3). Sequences similar or identical to eukaryotic NLSs are present in proteins in different prokaryotes (4–12), indicating that the origin of NLSs precedes the separation of the nucleus and the cytoplasm. This integration of NLS(s) into different (mostly DNA- and/or RNA-binding) domains was also observed in modern Eukaryota (11, 13–16).

Separation of the genome from the cytoplasm was followed by the evolution of numerous biomolecular condensates with concentrated proteins and/or RNAs and compartmentalized intranuclear processes (17–20). The largest intranuclear condensate is the nucleolus, which integrates numerous functions for genome organization and function (21). Some proteins possess special signal sequences, which are referred to as nucleolar localization signals (NoLSs), to enable their accumulation inside nucleoli (22). The origin of the NoLS is unclear.

Viruses have adapted amazingly to infecting and replicating in eukaryotic cells by hijacking nuclear structures and processes (23). Viruses evolved long before the emergence of Eukaryota, when the unique ancestor of living organisms commonly referred to as the last universal cellular ancestor (LUCA)) had acquired a highly complex virome (24). In the course of evolution, viruses adapted to an increasingly complex environment of ancestral eukaryotic cells. The size of viral particles did not substantially increase during evolution (e.g., 50-90% of virus particles detected in marine virome have an average diameter of 50 nm (25)). Smaller viruses have multiple fitness advantages since a greater number of viral offspring can be created from limited host resources; smaller particles also diffuse to encounter new hosts more rapidly. Virion size and genome size are tightly correlated (26); therefore, proteins encoded by viral genomes are generally small and often versatile: оne viral protein may be involved in many different processes, including regulation of the viral life cycle and modulation of host cell functions. The integration of the NLS, and probably the NoLS, into different functional domains might confer special benefits to viruses by allowing them to express small proteins.

Here, we used the trans-activator of transcription (Tat) protein of human immunodeficiency virus 1 (HIV-1) to investigate how viral proteins integrate the NLS and NoLS domains. HIV-1 Tat is a regulatory protein essential for productive and processive transcription from the HIV-1 long terminal repeat (LTR) promoter. Tat binds to viral RNA and recruits a complex of cyclin T1 and cyclin-dependent kinase 9 (CDK9) that activates RNA polymerase II, thus increasing transcription from the viral promoter (27). Tat is a small protein encoded by two exons (28) containing several domains: (i) the N-terminal proline-rich domain (1-21 aa), (ii) cysteine-rich domain (22-37 aa), (iii) hydrophobic core domain (38-47 aa), (iv) basic domain (BD) (48-59 aa), (v) glutamine-rich domain (60-72 aa), and (vi) C-terminal domain. While Tat can tolerate sequence mutations as high as 38% without its activity significantly changing (29), the BD is highly conserved in Tat variants. It is enriched with positively charged arginine and lysine residues comprising the ^49^RKKRRQRRR^57^ motif. The BD confers many properties to Tat (for a review, see (30)). To activate viral transcription, Tat binds *via* its BD to a short nascent stem-bulge loop leader RNA transactivation responsive region (TAR) at the 5’ extremity of viral transcripts (27, 31, 32). The BD also functions as an NLS (28, 33–36); however, the mechanism of Tat nuclear import is still debated. While some data indicate that Tat is imported into the nucleus *via* its interaction with importin-*α* (37–39), *in vitro* assays suggest that Tat nuclear import is mediated by the direct binding of its BD to importin-*β* (40). Additionally, the Tat BD interacts with nuclear components, probably with RNAs, which can also lead to nuclear accumulation (36). The BD also functions as an NoLS (28, 33, 41, 42); Tat accumulation might occur *via* an electrostatic interaction of its BD with nucleolar components (42). The accumulation of Tat inside nucleoli modulates nucleolar processes (43, 44).

Thus, the structure of the Tat BD allows the realization of several unrelated functions. This multifunctionality can be beneficial for the virus, as it acquired several functions without an increase in the protein size. Here, we investigated this multifunctionality of the Tat BD and demonstrated that NLS and NoLS were inevitably integrated into the HIV-1 Tat BD. Integration of an NLS and/or NoLS into functional protein domains might influence viral evolution, since this integration may lead to a reduction in protein size.

## RESULTS

### Tat is actively imported into the nucleus *via* its BD

To investigate the mechanisms of nuclear and nucleolar accumulation of the HIV-1 Tat protein in living cells, we used Tat protein fused with EGFP (EGFP-Tat). To exclude the possibility that EGFP inhibited Tat transactivation activity, we developed an *in vivo* assay based on a fast-maturing TurboRFP fluorescent protein controlled by a fragment of the HIV-1 3’ LTR that included the TAR (LTR-TurboRFP plasmid). We cotransfected U2OS cells with plasmids coding EGFP-Tat and LTR-TurboRFP and observed TurboRFP fluorescence in cells coexpressing EGFP-Tat but not EGFP or EGFP-BD (Fig. S1A). The results showed that the Tat protein after fusion with EGFP (EGFP-Tat) retained the ability to transactivate viral transcription. Additionally, we estimated the TurboRFP fluorescence intensity using flow cytometry and found that only EGFP-Tat had the transactivation capacity as compared to two mutant variants of Tat protein (EGFP-TatC22A, a mutant protein deprived of transactivation capacity because of 22C→A substitution, and EGFP-TatΔBD), EGFP and EGFP-BD (Fig. S1B).

We next analyzed the subcellular localization of EGFP-Tat transiently expressed in U2OS cells. EGFP-Tat was detected predominantly inside nuclei (Fig. 1A), where it accumulated inside nucleoli (Fig. S1C). Tat with the deleted BD (EGFP-TatΔBD) was diffusely distributed in cells, while BD-fused EGFP (EGFP-BD) accumulated inside nuclei and nucleoli (Fig. 1A). To estimate the efficiency of nuclear accumulation, the ratio of nucleoplasmic to cytoplasmic fluorescence (F_nuc_/F_cyt_) was measured after background correction, and the measurements confirmed that the BD was essential for nuclear localization of the Tat protein (Fig. 1B). Similarly, preferentially nuclear localization was detected in HIV-1-infected HEK cells after staining with anti-Tat antibodies (Fig. S1D).

**FIG 1.**
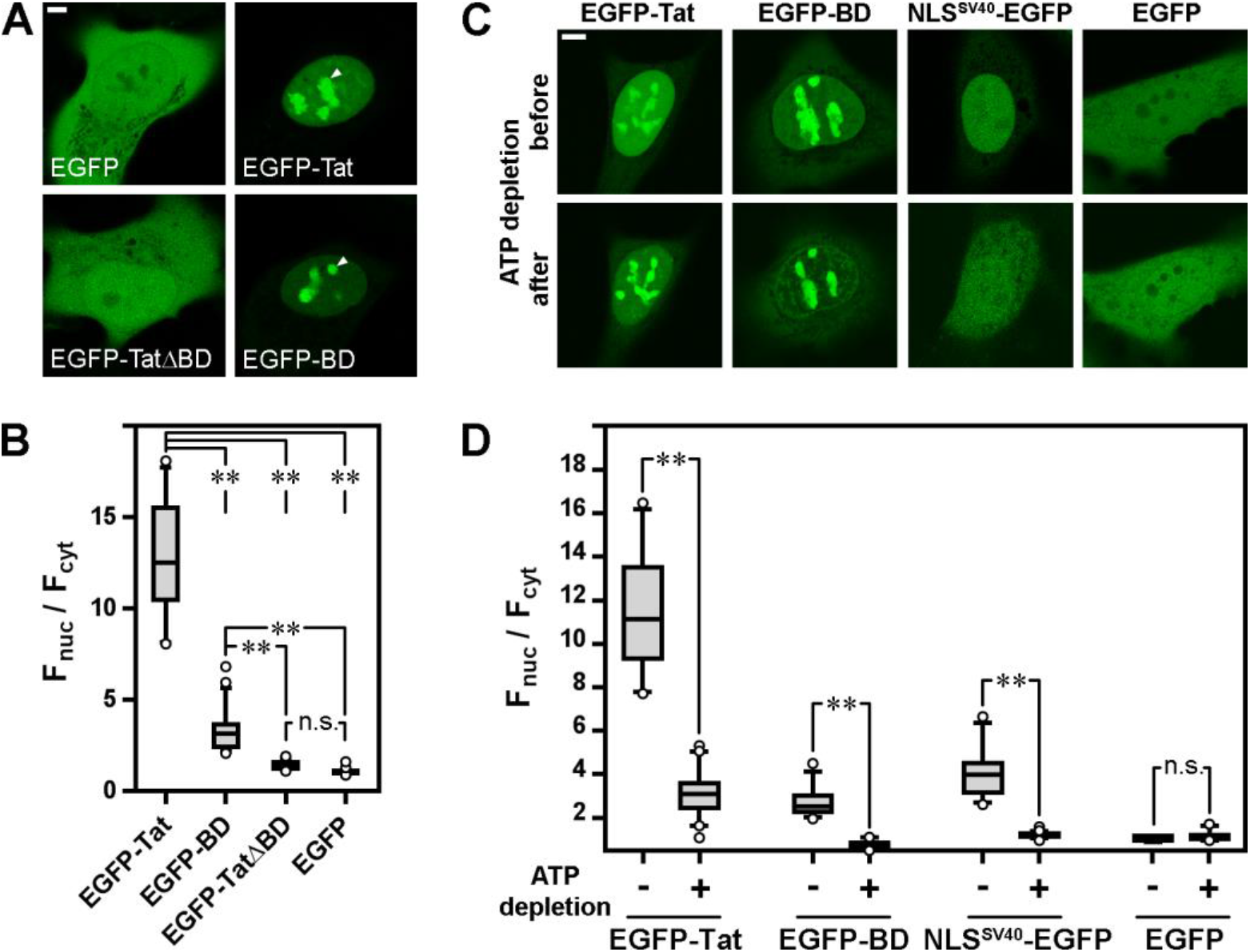
Basic domain of HIV-1 Tat functions as an NLS. (A) The localization of EGFP, EGFP-Tat, EGFP-TatΔBD and EGFP-BD in U2OS cells (live cell imaging). The intranuclear accumulation of EGFP-Tat and EGFP-BD (arrowheads) corresponded to nucleoli (see also Fig. S2). Bar – 5μm. (B) Estimation of the nuclear accumulation (Fnuc/Fcyt) of EGFP, EGFP-Tat, EGFP-TatΔBD and EGFP-BD in living U2OS cells. The comparisons were performed using the Kruskal-Wallis test (n.s. – not significant; ** – p < .005; n > 35). (C) The localization of EGFP-Tat and EGFP-TatΔBD in living U2OS cells before and after ATP depletion. EGFP-NLS^SV40^ was used as a positive control, and EGFP was used as a negative control. The cells were observed under identical conditions, and the images of untreated and treated cells were processed similarly. For image processing of cells expressing EGFP-Tat, the gamma value was set to 2.5. Bar – 5μm. (D) Estimation of the nuclear accumulation (Fnuc/Fcyt) of EGFP-Tat and EGFP-TatΔBD in living U2OS cells before and after ATP depletion. The comparisons were performed with Mann-Whitney U tests (n.s. – not significant; ** – p < .005, n > 35).

There are two major ways that a protein uses to enter the nucleus: passive diffusion, which is possible only for small proteins (<40-60 kDa), and active transport *via* importin-dependent pathways. The size of the EGFP-Tat fusion protein in this study was within the limit for passive diffusion, and Tat protein was shown to freely diffuse through nuclear pore complexes (45). However, diffusion *per se* cannot lead to nuclear accumulation: biologically inert EGFP freely enters the cell nucleus by diffusion, but its concentration is roughly the same in the nucleus and cytoplasm (Fig. 1A and B). Nuclear accumulation might result either from nuclear retention *via* interaction with nuclear components or from active nuclear import *via* importin-dependent pathways. Nuclear import is an energy-dependent process, and ATP depletion can be used to discriminate between nuclear retention and active nuclear import (46). For ATP depletion, U2OS cells expressing EGFP-Tat were incubated in a buffer containing sodium azide and 2-deoxyglucose for 50 min. Cells expressing either EGFP or SV40 T-antigen NLS fused with EGFP (NLS^SV40^-EGFP) were used as the negative and positive controls, respectively. Live-cell imaging demonstrated that NLS^SV40^-EGFP was relocalized from the nucleoplasm into the cytoplasm upon ATP depletion, while EGFP localization was not changed (Fig. 1C). We also observed a substantial decrease in the nucleoplasmic concentrations of EGFP-Tat and EGFP-BD in the absence of ATP (Fig. 1C and D). Nuclear accumulation of EGFP-Tat decreased after ATP depletion, but the level remained higher than that of EGFP and NLS^SV40^-EGFP under similar conditions, suggesting that nuclear accumulation of EGFP-Tat was partially due to active transport and, to a lesser extent, to nuclear retention. Importantly, after ATP depletion, the nuclear accumulation of EGFP-BD decreased to the level of NLS^SV40^-EGFP and EGFP, indicating that while active nuclear import was provided by the BD, other domains of the Tat protein were essential for its nuclear retention.

### Localization of the NLS in the Tat BD

The most common type of NLS, classical NLS (cNLS), is imported into the nucleus *via* interaction with importin-*α*. The most characterized monopartite cNLS has a consensus sequence of K(K/R)X(K/R), ensuring the optimal interaction with importin-*α* (2). The Tat BD contains nine amino acids (^49^RKKRRQRRR^57^) and thus is substantially longer than necessary for binding to importin-*α*. We used several software progrаms to predict the potential NLS(s) in the BD. NLStradamus (47), seqNLS (48), NucPred (49) and cNLS Mapper (50) predicted an NLS that roughly corresponded to the BD (Fig. 2A), while PSORT II (51) predicted three NLSs (^49^RKKR^52^, ^50^KKRR^53^ and ^55^RRRP^58^) overlapping the Tat BD. To map the NLS(s) in the BD, we systematically substituted each amino acid in the BD with alanine, followed by expression of the mutated Tat proteins and the analysis of their localization *in vivo* (Fig. 2B). The substitution of any arginine resulted in an ~2-fold decrease in Tat nuclear accumulation. The substitution of any lysine or glutamine with alanine decreased Tat accumulation, albeit to a lesser extent, and this difference was not statistically significant. Thus, arginine residues are more important than lysine residues for the NLS function of the Tat BD. These data seem to contradict the recently obtained crystal structure of the importin-*α*/TAT-NLS (^47^SGRKKRRQRRRAPQN^61^) complex, in which lysine residues constituted the main interaction core (39). To resolve this discrepancy, we employed molecular modeling.

**FIG 2.**
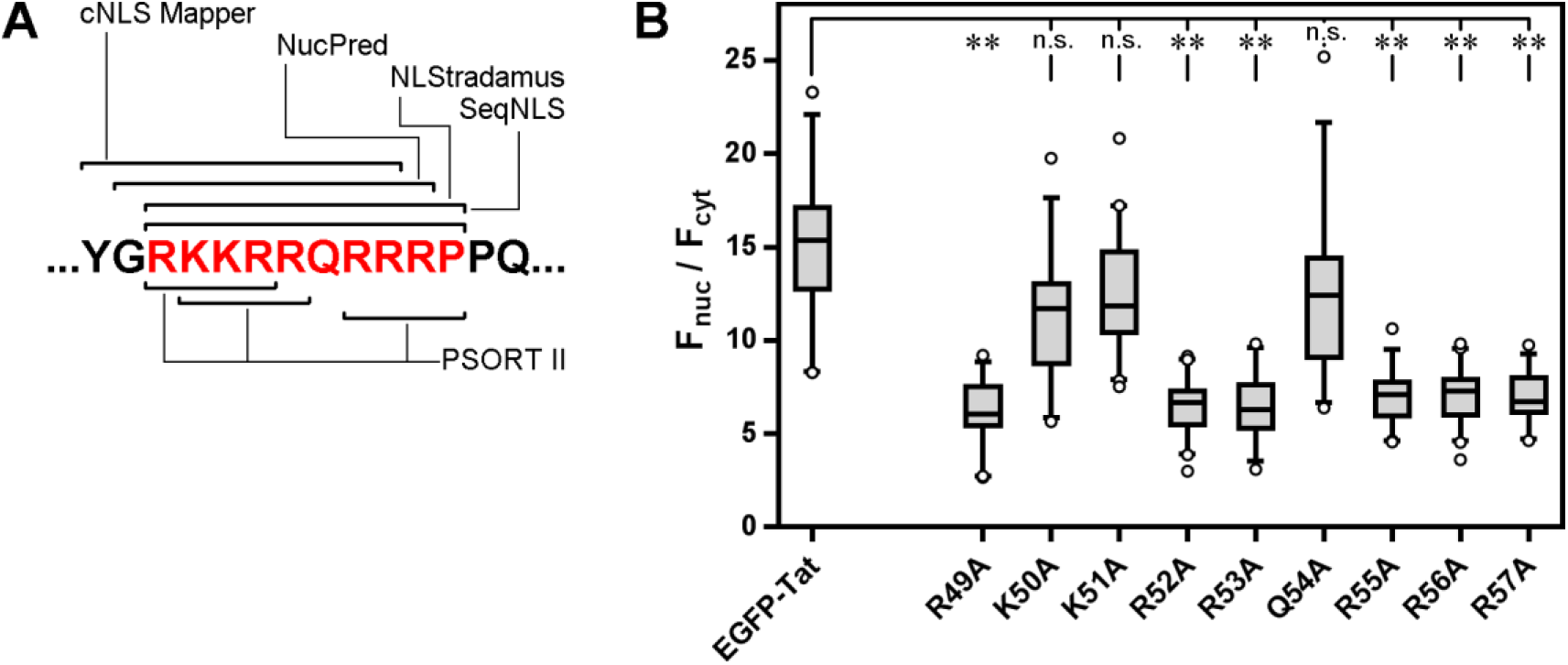
Prediction and mapping amino acids essential for nuclear accumulation of HIV-1 Tat. (A) Prediction of NLS in the BD of HIV-1 Tat by different programs: NLStradamus (2-state HMM static, prediction cutoff 0.6), seqNLS (final-score cutoff 0.86), NucPred (colored from yellow to red), cNLS Mapper (cutoff score 7.0), PSORT II. Amino acids of the BD are highlighted in red. (B) Nuclear accumulation (F_nuc_/F_cyt_) of EGFP-Tat and its mutants in living U2OS cells. The comparisons were performed with Kruskal-Wallis tests (n.s. – not significant; ** – p < .005; n > 30).

A broad range of computational approaches is routinely applied to study and design interactions between macromolecules with atomic precision (52); they proved useful in resolving the contradictory structural data in previous studies (53) and in situations with multiple potential binding modes (54). To predict the potential interaction sites of different parts of the BD and importin-*α*, we performed two series of docking procedures. We started with rapid coarse grain docking of the ^47^SGRKKRRQRRR^57^ peptide into the full importin-*α* structure. The docking experiment revealed two main binding sites overlapping the major and minor NLS-binding sites (Fig. 3A). Of the ten final models, two models showed binding outside the NLS-binding sites, six showed binding to the major NLS-binding sites, and two showed less favorable configurations with binding at the minor site, an interaction that was not apparent in the crystal structure.

**FIG 3.**
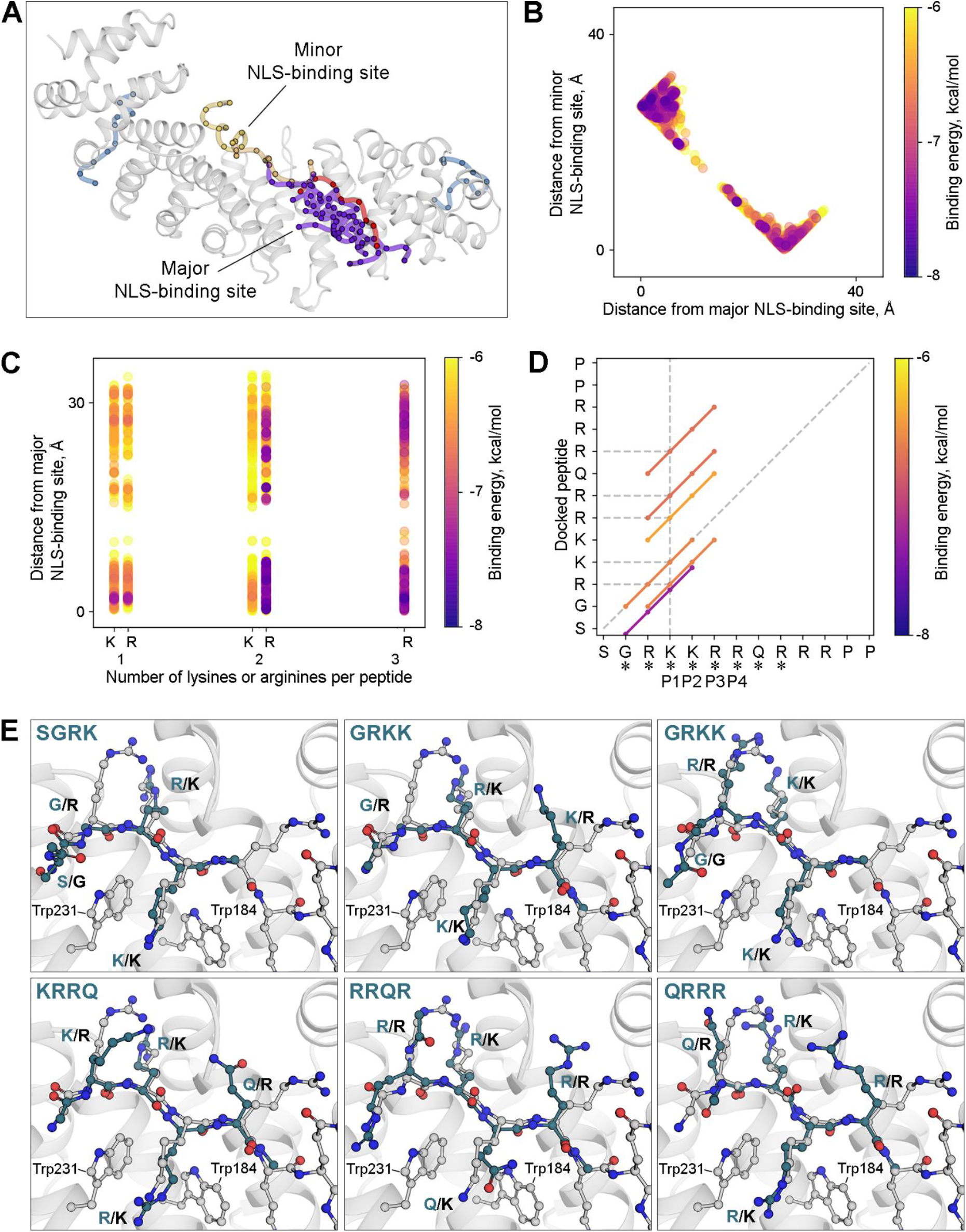
Molecular docking analysis of interaction of the Tat BD with importin-*α*. (A) Poses reported by CABS-dock. This approach utilizes a simplified representation of amino acids (each amino acid is represented by one coarse-grained particle). Each pose corresponds to a cluster of structures obtained during Monte Carlo modeling. Importin-*α* is depicted in gray, and the Tat NLS from the crystal structure is depicted in red. Major NLS poses are colored violet, minor NLSs are colored orange, and off-site structures are colored blue. (B) Distribution of docking energies between NLS binding sites obtained in the full-atom docking of the tetrapeptides. (C) A number of lysine residues or arginine residues per tetrapeptide. Estimated docking energies are lower (better) for peptides with more arginine residues. (D) Distribution of the best binding energies for tetrapeptides that comprise the NLS sequence. (E) The best binding poses of tetrapeptides of the NLS are in agreement with the published crystal structure. The published crystal structure is depicted in gray, and the tetrapeptides are depicted in green.

To obtain a detailed model of the interactions between the peptides and importin-*α* NLS-binding sites, we performed full-atom docking of tetrapeptides derived from the Tat NLS. In agreement with the coarse-grain docking, the major NLS-binding site attracted more peptides than other protein areas (Fig. 3B). Moreover, peptides with higher binding energy estimates were enriched at the major NLS-binding site, not the minor site (Fig. 3B). We next analyzed tetrapeptides in the major binding site and found that the arginine-rich tetrapeptides had a higher binding energy than the lysine-rich tetrapeptides (Fig. 3C). Importantly, we found that the arginine-rich peptides in different parts of the BD docked to the major NLS-binding site overlapping the GRKKR fragment of the X-ray structure (Fig. 3D). Although in case of some peptides, arginines occupied pockets initially assigned for lysines, the positions of peptide backbones were similar to the backbone position in the crystal structure (with the full-atom backbone RMSD less than 1 Å) (Fig. 3E). Despite the similarities between models with lysine and arginine, the difference in binding energies may result from an unusual amino acid composition of the binding sites. The major importin-α NLS binding sites were enriched in aromatic amino acids, especially tryptophan residues (Trp184, Trp 231 and Trp273), which possessed electron-rich conjugated pi-systems (Fig. 3E). In this context, recognition of positively charged organic cations is realized through electrostatic interactions. Surprisingly, arginine residues exhibit a stronger binding than lysine residues in pi-cation interactions (55), in agreement with the results of our mutagenesis study (Fig. 2B).

As we noted above, the analysis of docking results demonstrated that regardless of the peptide sequences, the backbone position was almost constant in most of the docking poses (Fig. 3E). This diversity of sidechain positions with the almost immutable backbone position in the crystal structure could be misinterpreted as a result of thermal fluctuations of atoms around their respective positions, thus causing an increase in the so-called thermal or B-factor. For instance, B-factors greater than 100 Å² or residues totally missing from the structure because of large thermal motions in the otherwise stable and well-resolved areas may indicate this type of problem region or be a sign of structure overfitting (53, 56, 57). The polder map around the NLS shows excessive unattributed electron density around N-terminal residues (Fig. S2). Moreover, many of the NLS residues have high B-factors, with that of the terminal arginine exceeding 100 Å². Thus, the reliability of the positions of these residues is relatively low.

The docking results indicated that different parts of the BD could interact with the major site of importin-*α*. To test whether the reported electron density map (39) can accommodate different peptides, we rescored the docking results in accordance with how well their heavy atoms (with the exception of hydrogen) fit in areas with higher levels of electron density (Fig. S3). Two peptides, SGRK and GRKK, had the highest enrichment among the top-scoring peptides. With 58 and 22 % of occurrence in the top-3 list from every run, they greatly outperformed the other peptides. Nevertheless, RKKR, KKRR, RQRR, and KRRQ were also able to adequately fit the electron density, although KKRR had the reverse orientation (Fig. S3). In the selected structures, arginine residues occupied sites initially assigned to N-terminal glycine or both lysine residues in the crystal reported structure. We observed multiple GRKK configurations, with one being identical to the crystal structure, while in the other, glycine and arginine interchanged their positions. Considering the unexplained residual electron density peaks at the −6 to +7 sigma level over G48 and R49 and a Ramachandran plot outlier at R49, we performed a correction of the initial 5SVZ model by rotating the G48 and R49 side chains about the R49 CA-C bond by 180 degrees following refinement with Refmac (58), resulting in R=0.16, Rfree=0.19 at 2.0 Å with no significant residual density over the remodeled segment (Fig. S4). R49 was in the preferred Ramachandran region, while its guanidine group established hydrogen bonds with side chains 234, 235, 270, and 277.

Hence, the peptide reported by (39) and several other peptides composing the Tat BD can be accommodated within the published crystal structure (Fig. 3E, Fig. S3).

The docking results indicated that the same positions could be occupied by either lysine or arginine. To ascertain whether this interchangeability between lysine residues and arginine residues is a unique feature of HIV-1 Tat, we analyzed all published PBD structures obtained for the complexes of importin-*α* with 38 eukaryotic and 14 viral proteins (Table S1). Monopartite cNLSs have a consensus sequence of K(K/R)X(K/R)(2). Consecutive residues from the N-terminal lysine of the monopartite NLS are referred to as P1, P2, etc. Previous structural (59, 60) and thermodynamic (61) studies demonstrated that a monopartite cNLS requires a lysine in the P1 position, followed by basic residues in positions P2 and P4. However, in three importin-*α*/NLS PDB structures obtained for viral proteins (beak and feather disease virus capsid protein (PDB ID 4HTV), HIV-1 viral protein R (Vpr) (PDB ID 5B56), and influenza A virus nucleoprotein (NP) (PDB ID 5V5O)), an arginine residue was substituted for a lysine in the P1 position (Fig. 4A; Fig. S5). We obtained a consensus logo structure for the region that directly interacted with importin-*α*. The P1 position was always occupied by lysine in eukaryotic proteins (Fig 4B), while the presence of arginine residues in the P1 position of some viral NLSs produced a slightly different consensus sequence of the viral NLS ((K/R)(K/R)X(K/R)) (Fig. 4C).

**FIG 4.**
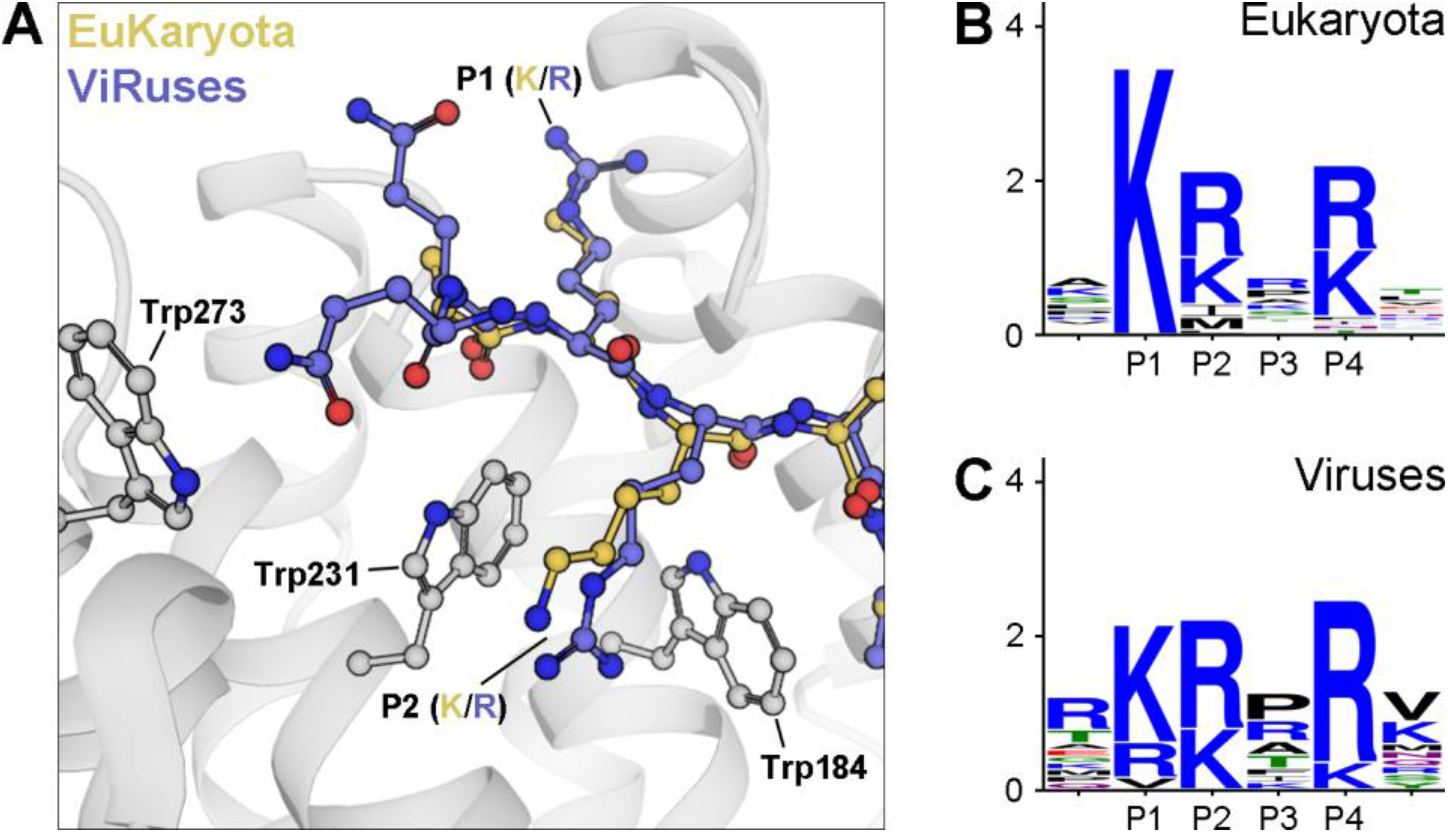
Differences between the eukaryotic and viral NLS consensus sequences. (A) Interaction of a typical NLS in a eukaryotic protein (Ku80, PDB ID 3RZ9) and a viral protein (HIV-1 VPR, PDB ID 5B56) with importin-*α*. (B) LOGO sequence of the region directly interacting with importin-*α* obtained from 38 PBD structures formed by importin-*α* and eukaryotic NLSs (Table S1). Consecutive residues from the N-terminal highly conserved lysine residue are referred to as P1, P2, etc. (C) The LOGO sequence of the region directly interacting with importin-*α* was obtained from 14 PDB structures formed by importin-*α* and viral NLSs (Table S1). In viral NLSs, the P1 position, which is occupied only by lysine in eukaryotic NLSs, can contain an arginine or a valine residue.

Hence, NLSs from eukaryotic proteins always (at least in all described cases) contain lysine in the P1 position in the major NLS-binding site, indicating the functional importance of this lysine; thus, the NLS sequence might be suboptimal for nuclear import in some viral proteins.

### Long NLSs in viral proteins

To ascertain whether similar long regions with NLS-like activity are present in other viral proteins, we analyzed published data on experimentally reported viral NLSs in which the NLSs were precisely mapped using either site-directed mutagenesis or importin binding. We found 10 viral proteins with either long (≥9 a.a.) or adjacent NLSs (Table S2). Some of these “long” NLSs, as in the case of NLSs in the feline immunodeficiency virus Rev protein, contained a long region enriched with positively charged amino acids similar to the Tat BD (62); others, e.g., 2b protein in cucumber mosaic virus, contained two or more adjacent NLSs (^22^KKQRRR^27^ and ^33^RRER^36^) (63). Thus, viral proteins may contain extended regions that function as NLSs.

The short classical NLSs and long NLSs described above represent two extreme variants. To analyze all variants and describe the heterogeneity of NLS structure, we assessed 106 NLSs in 88 viral proteins and 269 NLSs in 228 human proteins (Tables S3). All these NLSs, both in viral and human proteins, were enriched with positively charged amino acids (arginine residues and lysine residues)(Fig. 5A). Importantly, viral NLSs contained more arginine residues than lysine residues, while human NLSs were lysine-rich. Surprisingly, the prevalence of arginine residues over lysine residues is an overall characteristic of NLS-containing viral proteins; in contrast, the prevalence of lysine residues over arginine residues is seen in NLS-containing human proteins.

**FIG 5.**
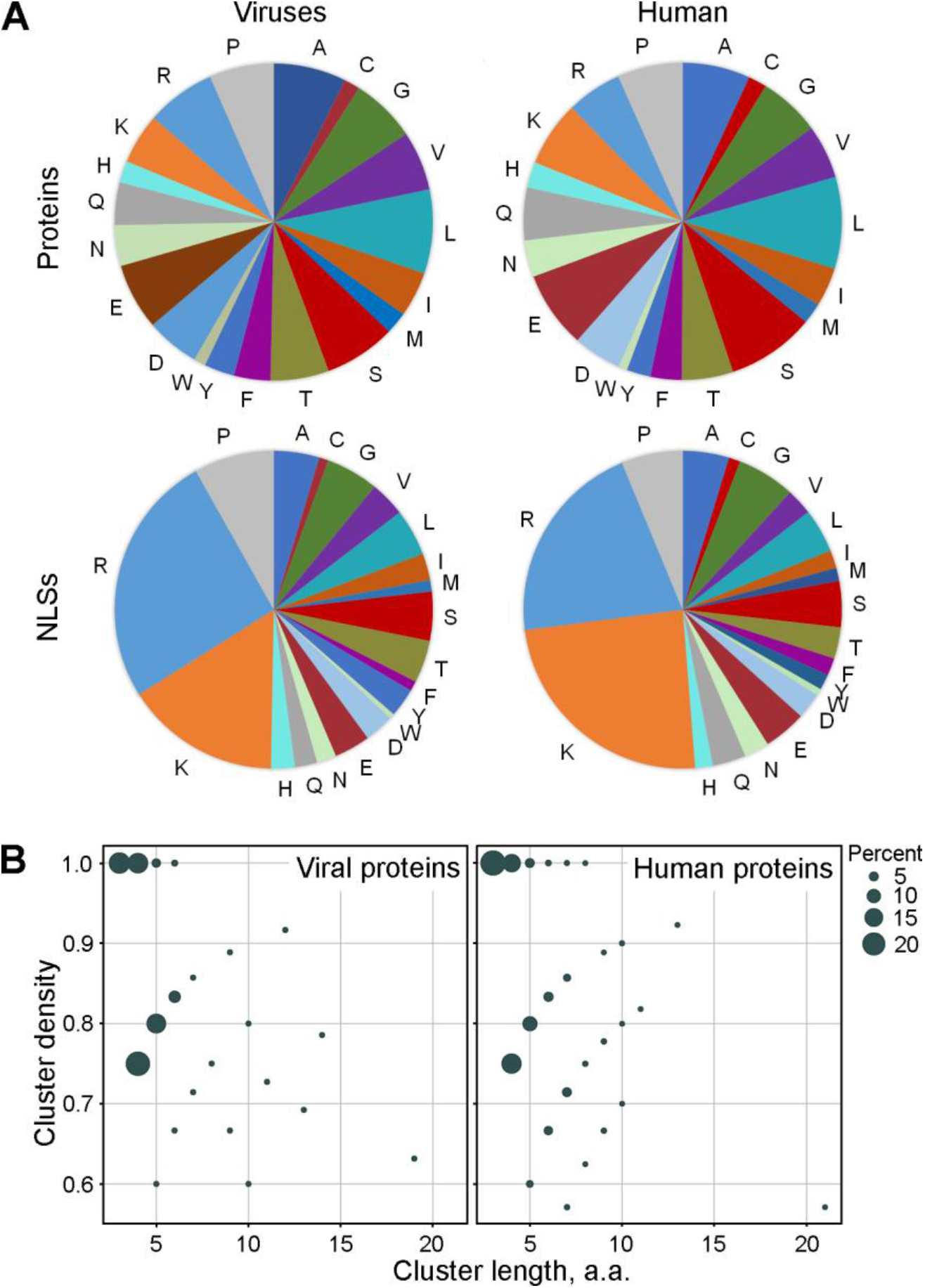
Heterogeneity of NLS organization in viral and human proteins. (A) The average amino acid content of known NLSs in viral and human proteins (bottom panels) in comparison with the overall amino acid content of these proteins (top panels). (B) Size of clusters of positively charged amino acids within human and viral experimentally confirmed NLSs. The clusters are defined as sequential arginine residues/lysine residues/histidine residues that are interspersed with no more than one non-positively charged amino acid.

The NLSs were analyzed for the size and structure of clusters of positively charged amino acids (Fig. 5B). We identified all clusters where arginine residues/lysine residues/histidine residues were interspersed with no more than one non-positively charged amino acid. The clusters containing three and four positively charged amino acids corresponded to cNLSs (K(R/K)X(R/K)). The number of these clusters in viral NLSs was slightly higher than that in human NLSs (~39% of viral clusters and ~32% of human clusters), while the number of clusters with 7-10 amino acids was twofold greater in human NLSs (~15%) than in viral NLSs (~6%), probably due to the presence of additional types of noncanonical NLSs that seem to be rare in viral proteins. Hence, long NLSs are present in both viral and human proteins.

### Tat accumulation inside nucleoli depends on the charge of its BD

Previous studies showed that Tat protein accumulated inside nucleoli and that this accumulation depended on the BD (28, 33, 41, 42). We confirmed these observations by expressing Tat-EGFP and its mutant forms in U2OS cells (Fig. 6A). Deletion of the BD led to a reduction in Tat-EGFP nucleolar accumulation, and the BD alone led to the accumulation EGFP in nucleoli, indicating that the BD indeed functioned as an NoLS. While ATP depletion led to a decrease in nuclear accumulation (Fig. 1C and D), the nucleolar accumulation of Tat simultaneously increased (Fig. 6B). We quantified the nucleolar accumulation by measuring the ratio of EGFP fluorescence in the nucleolus to that in the nucleoplasm (F_no_/F_nuc_) and found that nucleolar accumulation was ~2-fold higher after ATP depletion. NoLS(s) facilitate protein accumulation inside nucleoli by interacting with nucleolar components (nucleolar retention) (42, 64). To ascertain whether Tat nucleolar retention changed after ATP depletion, we analyzed the EGFP-Tat exchange between the nucleolus and the nucleoplasm using FRAP (Fig. 6C). In untreated U2OS cells, full recovery of EGFP-Tat was observed within t1/2 ~6 s. Upon depletion of cellular ATP, the recovery rate was decreased to ~65%, indicating that a substantial fraction of EGFP-Tat was immobilized inside nucleoli after ATP depletion (Fig. 6D). The t1/2 for the mobile fraction was ~9 s, and thus, the rate of Tat exchange was decreased. Thus, the retention of the Tat protein inside nucleoli led to nucleolar accumulation.

**FIG 6.**
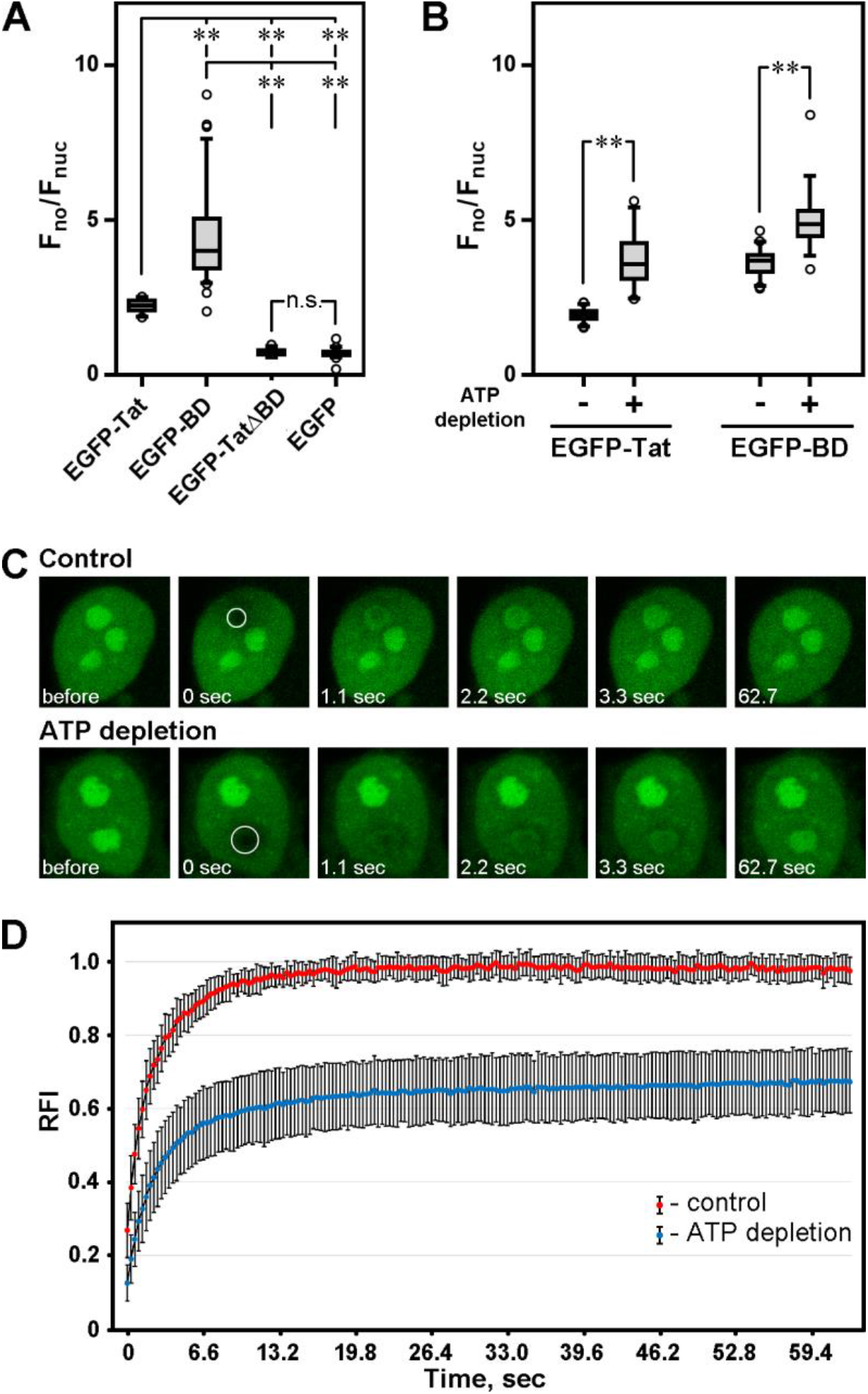
Nucleolar accumulation of HIV-1 Tat depends on its BD. (A) Nucleolar accumulation (F_no_/F_nuc_) of EGFP, EGFP-Tat, EGFP-TatΔBD and EGFP-BD in living U2OS cells. The comparisons were performed with Kruskal-Wallis tests (n.s. – not significant; ** – p < .005; n > 20). (B) Nucleolar accumulation (F_no_/F_nuc_) of EGFP-Tat and EGFP-TatΔBD in U2OS cells before and after ATP depletion. EGFP-NLS^SV40^ was used as a positive control, and EGFP was used as a negative control. The comparisons were performed with Mann–Whitney U tests (n.s. – not significant; ** – p < .005; n > 35). (C) FRAP analysis of the EGFP-Tat interaction with the nucleoli in living U2OS cells. Cells expressing EGFP-Tat were imaged before and after photobleaching (the bleached regions are outlined). The contrast was normalized to adjust for the loss of fluorescence during imaging. (D) FRAP analysis of EGFP-Tat mobility in nucleoli of the control U2OS cells and after ATP depletion. The results are presented as the means±s.d. (n=28 and 38).

We next used site-directed mutagenesis to analyze the role of each amino acid in BD nucleolar accumulation. When any amino acid of the BD was substituted with alanine, nucleolar accumulation of EGFP-Tat decreased regardless of the residue substituted, with the exception of glutamine (Fig 7A). Tat nucleolar accumulation decreased further upon simultaneous substitution of two (Fig. 7B) or three (Fig. 7C) amino acids with alanine residues. To compare the effect of the mutation of one, two or three amino acids on nucleolar accumulation, we combined the data on all mutants (with the exception of the Q54A mutant, which did not affect nucleolar accumulation) with the same number of substitutions (Fig. 7D).

**FIG 7.**
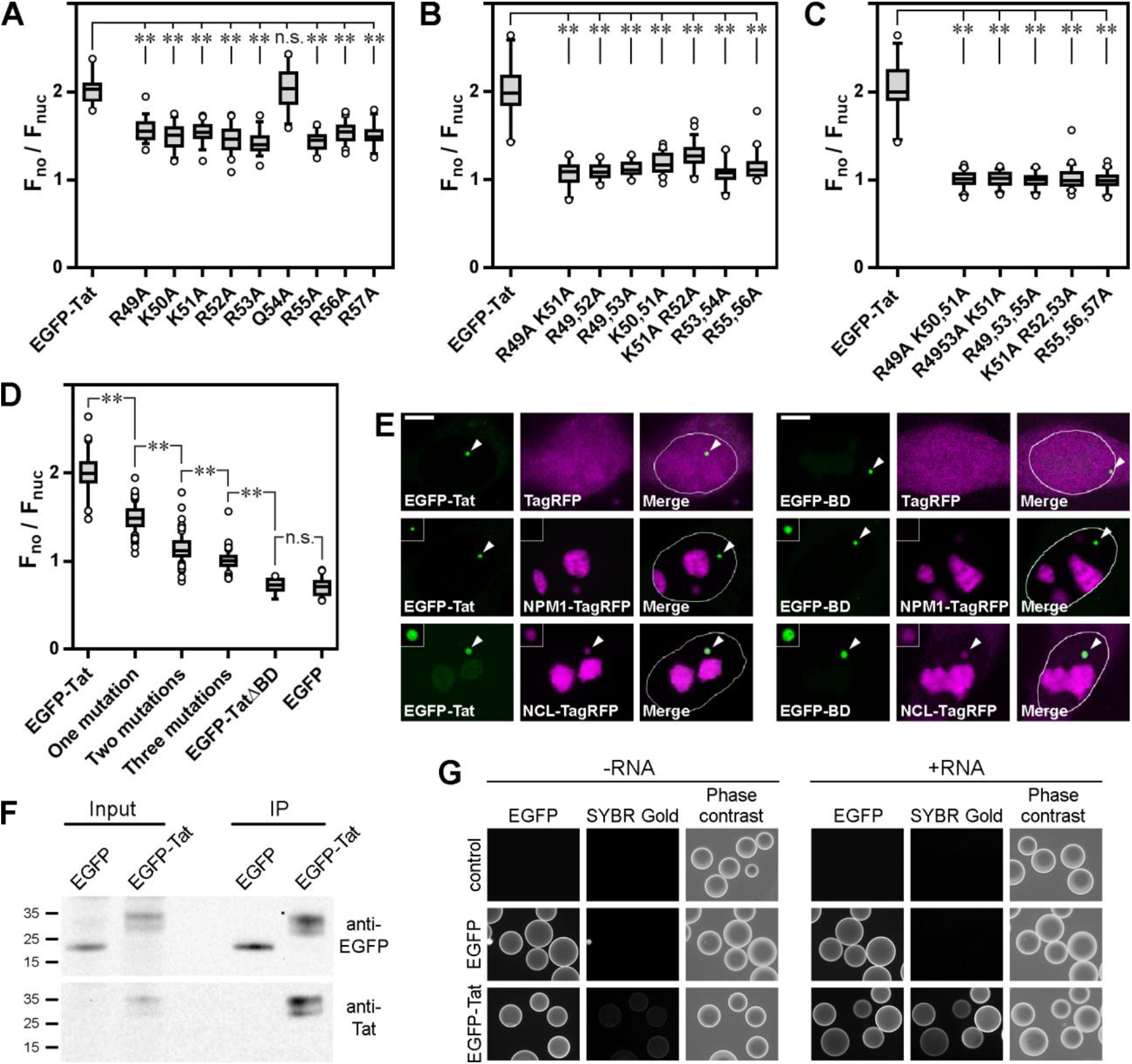
HIV-1 Tat BD interacts with nucleolar components. (A) Nucleolar accumulation (F_no_/F_nuc_) of the EGFP-Tat protein and its mutants (one a.a. substitution in the BD) in living U2OS cells. The comparisons were all performed with Kruskal-Wallis tests (n.s. – not significant; ** – p < .005; n > 20). (B) Nucleolar accumulation (Fno/Fnuc) of the EGFP-Tat protein and its mutants (two a.a. substitutions in the BD). (C) Nucleolar accumulation (F_no_/F_nuc_) of the EGFP-Tat protein and its mutants (three a.a. substitutions in the BD). (D) Nucleolar accumulation (Fno/Fnuc) of EGFP-Tat and combined data for all mutants (with the exception of Q54A) with an equal number of substitutions (i.e., one, two or three substitutions irrespective of the precise position of substitution). EGFP-TatΔBD and EGFP were used as negative controls. (E) *In vivo* interaction of HIV-1 Tat and its BD with NPM1 and NCL. The expressed EGFP-fused proteins were immobilized on a platform inside F2H cells (inserts), and the colocalization of the coexpressed proteins fused with TagRFP indicates the interaction between the two proteins. The contrast and brightness were adjusted for display purposes. For image processing of the cells expressing NCL-TagRFP, the gamma value was set to 2.0. (F) Western blotting of the cell lysates (input) and proteins immunoprecipitated on agar beads (IP) using antibodies against EGFP and the Tat protein. EGFP and EGFP-Tat were immobilized for a subsequent study on their interactions with cellular RNA. (G) Interaction of RNA with the Tat protein immobilized on agar beads. EGFP or EGFP-Tat proteins expressed in U2OS cells were immunoprecipitated on agar beads and then incubated with or without total cellular RNA from U2OS cells (+RNA or -RNA samples, respectively). The cells were observed and the images processed under identical conditions.

We also mutated each positively-charged amino acid within the BD to another positively-charged amino acid (i.e., R→K or K→R), assuming that such substitution should not substantially influence nucleolar accumulation if the accumulation was charge-dependent. Indeed, these substitutions did not affect the nucleolar accumulation of Tat, in agreement with our assumption (Fig. S5). Thus, nucleolar accumulation was indeed dependent on the proportion of positively charged amino acids, indicating that HIV-1 Tat accumulated inside nucleoli *via* a charge-dependent, and not a sequence-dependent mechanism.

### HIV-1 Tat interacts with nucleolin and nuclear RNA

The accumulation of proteins inside nucleoli was previously shown to result from high-affinity binding interactions with core nucleolar components, including ribosomal DNA, RNAs and proteins (65). Tat nuclear interactome includes several nucleolar proteins, including NPM1 and NCL (37), indirectly indicating that Tat can accumulate inside nucleoli due to its interactions with nucleolar proteins. The interaction of Tat with NPM1 was also previously demonstrated *in vitro* (66). However, NoLS(s) were shown to rather interact with nucleolar RNAs (42, 64). To test the potential interactions of the Tat protein with major nucleolar proteins *in vivo*, we used a fluorescent two-hybrid (F2H) assay to make use of cells with a GFP-anchoring platform in the nucleus. EGFP-fused proteins were tethered to the intranuclear GFP-anchoring platform and then assayed for colocalization with TagRFP-fused proteins that could potentially interact with GFP-fused proteins. After expression in F2H cells, EGFP-Tat was located preferentially at the GFP-anchoring platform in the majority of cells (nucleolar accumulation was clearly visible only in cells with a high expression of EGFP-Tat, indicating a robust interaction with the platform) (Fig. 7E). We next analyzed the interaction of EGFP-Tat with two major nucleolar proteins, NPM1 and NCL. After coexpression of EGFP-Tat and NPM1-TagRFP, we did not observe any accumulation of NPM1-TagRFP at the platform. In contrast, clear colocalization at the platform was seen after coexpression of EGFP-Tat and NCL-TagRFP, indicating a direct interaction between these proteins. Interestingly, some accumulation of NCL-TagRFP on the platform was detected in ~50% of the cells after EGFP-BD and NCL-TagRFP were coexpressed, indicating that the interaction between Tat and NCL can occur through the BD (Fig. 7E).

We next analyzed whether Tat interacted with cellular RNAs using an RNA pull-down assay. EGFP-Tat or EGFP was expressed in U2OS cells and then immunoprecipitated with magnetic beads (Fig. 7F). The beads were then incubated with total cellular RNA from U2OS cells and stained with SYBR-Gold dye. Fluorescence microscopy revealed that immunoprecipitated EGFP-Tat immobilized RNA on the surface of the agarose beads, while RNA was not detected on the surface of either the control beads or beads with EGFP (Fig. 7G); therefore, Tat protein interacts with both NCL and RNA, which may account for its retention in the nucleus and nucleolus.

### Overlapping of NLSs and NoLSs

Several reports previously described overlapping NLS(s) and NoLS(s) (for discussion, see (67)); however, a comprehensive mapping of both NLSs and NoLSs has never been performed. We used the datasets of experimentally established NLSs (Tables S3) and predicted NoLSs (68). In addition, we used a dataset of experimentally established NoLSs for which we predicted NLSs using the NLStradamus program (Table S4). Both prediction strategies demonstrated that NLSs and NoLSs more frequently overlapped in viral proteins than in human proteins (Fig. 8A).

**FIG 8.**
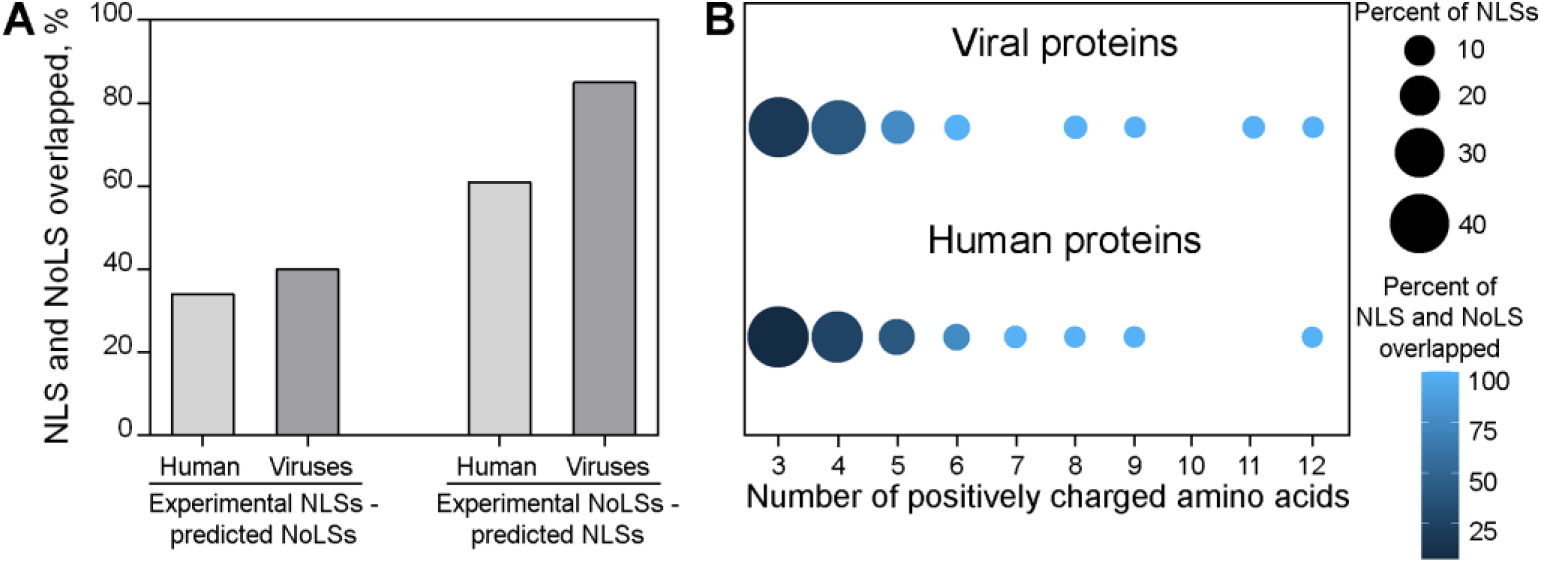
NLSs overlap with NoLSs in viral and human proteins. (A) Overlapping of experimentally annotated NLSs with predicted NoLSs and predicted NLSs with experimental NoLSs. (B) Long NLSs more frequently overlapped with the predicted NoLSs.

Estimates of the minimal content required for NoLS function vary. According to one report, a protein region can function as an NoLS if it contains more than either three arginine residues or five lysine residues (42), according to the other report, six arginine residues are required (64). NLSs are enriched with positively charged amino acids, and therefore, one can assume that long NLSs may function as NoLSs. Indeed, the majority of the long NLSs (with >6-7 positively charged amino acids) were predicted to be potential NoLSs (Fig. 8B).

## DISCUSSION

Many functions of HIV-1 Tat are attributed to its BD (30), the major function being the transactivation of viral transcription (69–72). Additionally, the Tat BD functions as an NLS and NoLS. Here, we investigated the mechanisms of NLS and NoLS integration into the BD of HIV-1 Tat and analyzed the implications of this integration for virus evolution.

### Integration of NLS into the Tat BD

The Tat protein is small, and therefore, it was previously shown to diffuse freely through nuclear pore complexes (45). However, the presence of NLS led to a robust nuclear accumulation of HIV-1 Tat compared to biologically inert EGFP. A striking feature of the Tat BD is its length (9 amino acids), which is substantially longer than the minimum required for association with importin-*α* (4 amino acids). We carried out systematic site-directed mutagenesis of all amino acids of the Tat BD and found that the whole BD sequence functions as an NLS. *In silico* molecular docking experiments indicated that different regions of the BD might potentially interact with importin-*α*. Following these results, we proposed a model of multiple concurrent binding modes of the Tat NLS, in which different parts of the BD can interact with importin-*α* (Fig. 9A). It appears that a single NLS can bind with importin-*α* in a variety of ways because of the excess of charged amino acids, preserving the overall geometry without any significant decrease in binding energy. As a result, the nuclear accumulation can be a consequence of cumulative binding between different fragments of BD and importin-*α*.

**FIG 9.**
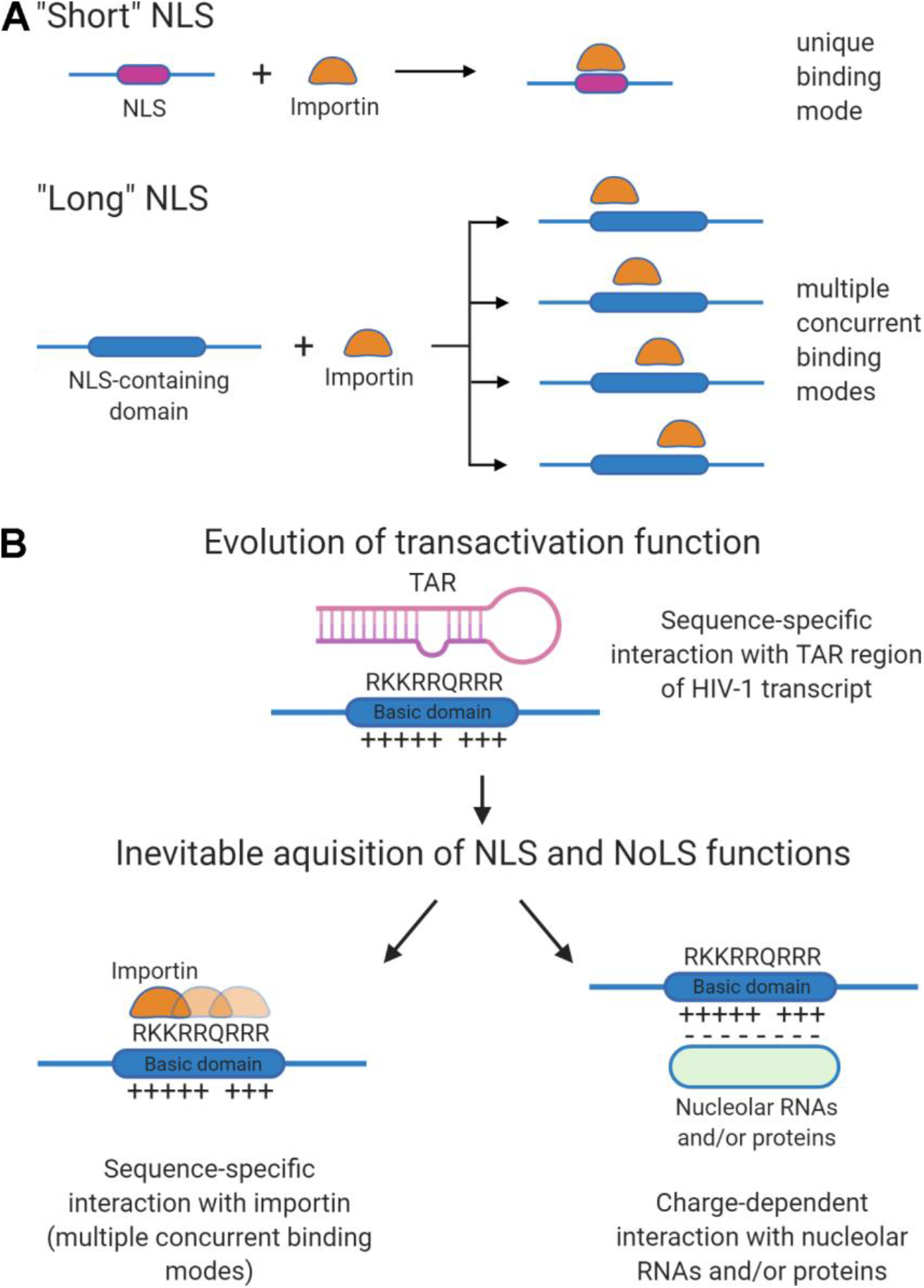
(A) Different regions of NLS-containing domains enriched with positively charged amino acids (“long” NLSs) might interact with importin-*α* (multiple concurrent binding modes), and nuclear accumulation may be a consequence of cumulative binding between different fragments of these regions and importin-*α*. (B) Evolution of domains enriched with positively charged amino acids may lead to the inevitable acquisition of additional functions (as an NLS and NoLS, as in the case of HIV-1 Tat), i.e., different functions evolve simultaneously as a complex of integrated activities (molecular coevolution).

The second unusual feature of the Tat NLS of HIV-1 is the essential role of arginine residues in NLS function, which was corroborated both by the results of site-directed mutagenesis and molecular docking. These observations were in obvious contradiction with the known consensus sequence of the classical monopartite NLSs, in which the P1 position is always occupied by lysine (K(R/K)X(R/K)). However, when we analyzed PDB structures obtained for complexes of importin-*α* with 38 eukaryotic and 14 viral NLSs, we found that in three NLSs from viral proteins, the most conserved lysine in the P1 position was replaced with arginine, and in one NLS, it was replaced with valine. Thus, at least in some viral proteins, NLSs have a suboptimal organization. Notably, the Tat BD contains a sequence that corresponds to the classical eukaryotic NLS consensus sequence (^50^KKRR^53^), but paradoxically, this region of the Tat BD did not lead to more efficient nuclear accumulation than other sequences in the BD. Thus, in some viral proteins, the NLS sequence is not optimal for nuclear import due to enrichment with arginine residues. This might be a consequence of the integration of NLSs into functional domains, as in the case of HIV-1 Tat. In the BD, arginine residues participate in binding to the TAR, which leads to transactivation of HIV-1 transcription (73, 74). This may be a general feature of these integrated NLSs, since 72-87% of known protein-RNA interfaces contain arginine residues (75, 76) whose substitution with lysine residues often diminishes their RNA-binding activity (73, 77, 78). Thus, when an RNA-binding domain acquires the function of an NLS, its RNA-binding and importin-binding efficiencies must be equilibrated. Moreover, we found that NLS-containing viral proteins are overall arginine-rich, and the composition of the viral NLS may simply reflect this enrichment.

Thus, integration of NLSs into functional domains may prevent the evolution of structurally optimal NLS sequences when evolution would impair the major function(s) of the domain, as in the case of HIV-1 Tat, but this might be compensated, for example, by the acquisition of multiple concurrent binding-mode mechanisms.

It appears that HIV-1 Tat is not a single viral protein with an integrated NLSs. The domains involved in RNA binding and the NLS of Herpes simplex virus type 1 nucleocytoplasmic shuttling protein UL47 overlap, and it is not possible to separate their activities (79). The organization of the NLS we found in Tat BD might be widespread not only among viruses but also among eukaryotes. Indeed, in some eukaryotic proteins, NLSs might also be integrated into other domains that occupy a significant portion of the protein. This situation was previously described for histones (80, 81), ribosomal proteins (82–84) and the nucleolar methyltransferase fibrillarin (FBL) (85). Histones and ribosomal proteins are small and highly conserved proteins, and the presence of extended NLSs may also be a consequence of their integration into a functional domain, which could not be uncoupled during evolution.

### Integration of NoLS into the Tat BD

NoLSs facilitate the accumulation of proteins inside nucleoli *via* a charge-dependent mechanism, not a sequence-dependent mechanism (42, 64, 86, 87); i.e., any positively charged region can potentially function as an NoLS. For example, core histones can accumulate inside nucleoli before their incorporation into chromatin or after their release from chromatin (86, 88–90), and it was demonstrated that histone H2B contains an NoLS in its N-terminal tail (86).

Here, we used site-directed mutagenesis to demonstrate that each positively charged amino acid of the BD is involved in the nucleolar accumulation of Tat; thus, Tat accumulated inside nucleoli *via* a charge-dependent mechanism, similar to other NoLSs. It seems logical that electrostatic interactions with nucleolar components that lead to nucleolar accumulation would have a low specificity. Indeed, we found that the Tat protein could interact with both the nucleolar protein NCL and RNAs. RNA is charged negatively, and NCL contains several extended acidic domains and even has a negative net charge at physiological pH (pI 4.1). Interestingly, we were unable to detect any interaction between Tat and NPM1, although it was previously demonstrated that Tat interacted *in vitro* with this nucleolar protein (37, 66).

Different functions integrated inside the BD may be regulated together. For example, Tat transactivation activity is affected by lysine acetylation (91–96) that regulates nuclear accumulation of proteins (97–101), and K51Q mutation of HIV-1 Tat which mimics acetylation, decreased the nuclear localization, indicating that lysine acetylation could modulate the subcellular localization of Tat, in addition to the regulation of its transactivation activity (94). As acetylation decreases the protein charge, it may also decrease the nucleolar accumulation. Thus, posttranslational modifications can simultaneously modify several functions within the Tat BD.

### Evolutionary implications of NLS and NoLS integration

Tat BD is a conserved region enriched with positively charged amino acids that serves as both an NLS and NoLS. The BD evolved primarily as a domain involved in binding with TAR and the transactivation of HIV-1 transcription, but its enrichment with positively charged amino acids inevitably led to the acquisition of its functions as an NLS and NoLS (Fig. 9B). These two additional functions are embedded in the BD amino acid sequence and cannot be uncoupled from the major function of the BD.

Evolution of the transactivation function would have been perturbed if nuclear and/or nucleolar accumulation had disrupted the performance of the main function of this BD. However, accumulation in the nucleus most likely promotes transactivation. Although the role of the nucleolar accumulation of Tat in HIV-1 infection was not directly demonstrated, it seems that due to the rapid exchange of Tat between the nucleoli and the nucleoplasm, its nuclear accumulation cannot negatively affect the transactivation function.

The primacy of the transactivation function during evolution led to the formation of NLSs which are substantially longer than classical NLSs. These NLSs might function through multiple concurrent binding with importin-*α*. Moreover, our bioinformatic analysis demonstrated that, in some viruses, the structure of the NLS may also be nonoptimal. The multiple concurrent binding mode mechanism allows viral proteins to accumulate in the nucleus due to the presence of several closely located or overlapping NLSs, even if a single NLS was not optimized for NLS function.

Examples of protein domain evolution when functions did not evolve sequentially and independently of each other but evolved directly in the form of an integrated functional complex (molecular coevolution) may be widespread, and traces of this integration might be observed in modern living beings. Indeed, NLS integration into annotated domains is a widespread phenomenon in Eukaryota (11). An additional indication of the inevitable character of NLS/NoLS function acquisition is the presence of these signals integrated into cytoplasmic proteins (16).

Integration of the NLS and NoLS into functional protein domains might influence viral evolution since this integration could prevent an increase in protein size. Separation of functions in different protein domains, which is fundamentally possible for eukaryotic proteins, leads to an increase in protein size that might be disadvantageous for viruses. Thus, the integration of the NLS and NoLS into functional domains could be a key phenomenon that influenced the origin and evolution of the NLS both in viruses and in eukaryotes.

## MATERIALS AND METHODS

### Cell culture

U2OS and HeLa cells (both from Russian Cell Culture Collection, Saint Petersburg, Russia) were grown in Dulbecco’s modified Eagle’s medium supplemented with alanyl glutamine (Paneco, Moscow, Russia), 10% fetal calf serum (HyClone, Logan, UT, USA) and an antibiotic and antimycotic solution (Gibco, Rockville, MD, USA). The HIV-1-virus was produced from 293T cells as previously described (102).

### *In vivo* analysis of transactivation activity of Tat protein

To detect the transactivation activity of the Tat protein, we developed an *in vivo* system based on the fast-maturing fluorescent protein TurboRFP. The fragment of the HIV-1 3’ LTR, which contains all of the U3 region and a fragment of the R region, including the TAR, was amplified from the HIV-1 LTR lacZ reporter vector, which was obtained through the NIH AIDS Reagent Program from Dr. Joseph Maio (103) using the following primers: 5’-AGTCAAGCTTTGGAAGGGCTAATTCACTCCCAAAG-3’ and 5’-AGTCGGTACCAGCTTTATTGAGGCTTAAGCAGTGGG-3’. The PCR product was digested with FastDigest HindIII and KpnI restriction enzymes (Thermo Scientific, Waltham, MA, USA) and cloned into a promoter-less vector encoding the red fluorescent protein TurboRFP (pTurboRFP-PRL; Evrogen, Moscow, Russia). U2OS cells were cotransfected with the EGFP-Tat, EGFP or EGFP-BD and LTR-TurboRFP plasmids, and the fluorescence was analyzed using an AxioVert 200M microscope (Carl Zeiss, Oberkochen, Germany) equipped with an ORCAII-ERG2 cooled CCD camera (Hamamatsu Photonics K.K., Hamamatsu City, Shizuoka, Japan).

### Plasmid construction

To construct EGFP-Tat, the pGST-Tat 1 86R plasmid was obtained through the NIH AIDS Reagent Program, Division of AIDS, NIAID, from Dr. Andrew Rice (104– 106). Full-length Tat was PCR amplified with Phusion high-fidelity DNA polymerase (Thermo Scientific, Waltham, MA, USA). The amplified PCR products were digested with BamHI and HindIII and inserted into a EGFP-C1 vector (Clontech, Mountain View, CA, USA).

For site-directed mutagenesis, the Tat gene was cloned into a pJET1.2/blunt cloning vector (Thermo Scientific, Waltham, MA, USA) using a CloneJET PCR cloning kit (Thermo Scientific, Waltham, MA, USA). Point mutations were made using a Change-IT multiple-mutation site-directed mutagenesis kit (Affymetrix, Santa Clara, CA, USA) according to the manufacturer’s instructions. Each subsequent round of mutagenesis was carried out after confirming the presence of the preceding necessary mutation by sequencing. The mutated genes were amplified by PCR; the resulting PCR products were digested by HindIII and BamHI, gel purified and cloned into a pEGFP-C1 vector (Clontech, Mountain View, CA, USA).

To construct the EGFP-BD plasmid, we designed adapters encoding the BD of HIV-1 Tat with HindIII and BamHI restriction sites. The oligonucleotides were incubated in annealing buffer (10 mM Tris, pH 8.0; 50 mM NaCl; and 10 mM MgCl2) for 5 min at 94°С and then for 5 min at 70°С. The resulting product was cloned into a pEGFP-C1 vector (Clontech, Mountain View, CA, USA).

To obtain the NCL-TagRFP plasmid, full-length NCL was PCR amplified with Phusion high-fidelity DNA polymerase (Thermo Scientific, Waltham, MA, USA) using the following primers: 5’-AGTCGAATTCATGGTGAAGCTCGCGAAGGCAG-3’ and 5’-AGTCGGATCCAATTCAAACTTCGTCTTCTTTCCTTGTGG-3’. The pEGFP-NCL expression plasmid (nucleolin), which was used as a template for PCR, was a kind gift from Dr J. Borowiec(107). The amplified PCR product was digested with EcoRI and BamHI and then inserted into the pTagRFP-N1 vector (Evrogen, Moscow, Russia). After cloning into an expression vector, the correctness of all constructs was confirmed by sequencing.

The construction of NLS^SV40^-EGFP (85) and NPM1-TagRFP (42) plasmids was previously described.

### Measurement of nuclear and nucleolar accumulation

To evaluate the nuclear and nucleolar accumulation of the Tat protein, we used a method described elsewhere with some modifications (42). Images of at least 20 living U2OS cells expressing EGFP-fusion proteins were acquired through two different experiments using a Nikon C2 confocal laser scanning microscope with a 63×1.4NA oil immersion objective under identical conditions. A region of interest was determined within the nucleolus, within the nucleoplasm and within the cytoplasm of the same cell. The mean gray value was determined for each, background levels were subtracted, and the nucleolar-nucleoplasmic (F_no_/F_nuc_) and nucleoplasmic-cytoplasmic (Fnuc/Fcyt) ratios were determined for every value pair. The statistical analysis and graph generation were performed using GraphPad Prism 6 software (GraphPad, San Diego, CA, USA). Statistical analyses were performed using GraphPad Prism 6 software using nonparametric Mann-Whitney U test or Kruskal-Wallis test with Dunn’s multiple-comparison test.

### ATP depletion assay

ATP depletion was carried out as described by (108). Briefly, HeLa cells were grown in 35-mm dishes on a coverslip (MatTek, Ashland, MA, USA). The cells were treated with 20% Hank’s solution for 15 min and then transferred into Medium 1 (150 mM NaCl, 5 mM KCl, 1 mM CaCl2, 1 mM MgCl2 and 20 mM HEPES, pH 7.4) containing 10 mM sodium azide and 6 mM 2-deoxy-D-glucose (Sigma-Aldrich, St. Louis, MO, USA) and incubated for 50 min. Live cell imaging was performed with a Nikon C2 confocal microscope with a 60× Plan-Apo objective (NA 1.4) at 37°C; focus stabilization was performed using a PFS system (Nikon, Minato City, Tokyo, Japan).

### Molecular docking

Docking was performed through two different approaches. Full-length coarse-grained docking of the SGRKKRQRRR peptides was performed with standalone CABS docking (109). The importin-*α* structure with PDB ID 5SVZ was cleared of the NLS peptide and water molecules. The default parameters for sampling efficacy (simulation cycles 50) were used, and no additional restraints were applied.

Full-atom docking of tetrapeptides was performed using the importin-*α* structure with PDB ID 5SVZ (a 30 Å x 45 Å x 45 Å docking box with the center at 80.559, 21.365, 100.184 was used to cover both NLS sites). Docking was performed with the Qvina-W package (110), which implements the AutoDock Vina (111) scoring function optimized for blind docking. Due to the relative complexity of peptide docking (112), the exhaustiveness parameter, which affects sampling efficacy, was set to 512. Twenty independent runs for every peptide were performed with twenty poses per peptide produced.

Rescoring of the poses was performed in accordance with a 2FoFc electron density map using a modified PeptoGrid (113) procedure. The density map was converted into a numerical grid with an elementary step of approximately 0.1 Å. Per-atom scores were calculated as negative log probabilities of corresponding grid cells with the corresponding sigma sign. Values less than +1σ were penalized as −3σ. Only heavy atoms were considered, with ACE and NME caps excluded from scoring. Total scores were sorted and normalized to the highest score.

### Fluorescence recovery after photobleaching (FRAP)

Cells were grown in 35-mm dishes on coverslips. The medium was overlaid with mineral oil before the experiment, and the dishes were mounted onto a Nikon A1 confocal microscope. Four single scans were acquired for the FRAP experiments, followed by a single pulse for photobleaching. The recovery curves were generated from background-subtracted images. The relative fluorescence intensity (RFI) was calculated as

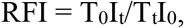

where *T_0_* is the total cellular intensity during prebleaching, *T_t_* is the total cellular intensity at time point *t*, *I_0_* is the average intensity in the region of interest during prebleaching, and *I_t_* is the average intensity in the region of interest at time point *t*. The results for at least 20 cells were averaged to obtain the final curve of fluorescence recovery.

### F2H assay

F2H cells (genetically modified baby hamster kidney (BHK) fibroblasts) were cultured according to the manufacturer’s instructions (ChromoTek GmbH, Planegg-Martinsried, Germany). For detection of protein-protein interactions in live cells, we transfected F2H cells with a plasmid coding the protein fused with EGFP and a second plasmid coding the protein fused with TagRFP. Cells were fixed with 3.7% paraformaldehyde, and images were acquired using a Nikon C2 confocal laser scanning microscope with a 63×1.4NA oil immersion objective.

### RNA pull-down assay

Total RNA was isolated from U2OS cells using an RNeasy mini kit (Qiagen Inc., Valencia, CA, USA) according to the manufacturer’s instructions. EGFP-Tat or EGFP genes were cloned into a pLCMV lentiviral vector. Lentiviral constructs were introduced into 293T cells along with packaging plasmids. Viral particles were used for the transduction of the U2OS cells, and cells expressing target proteins were selected using a FACSAria III cell sorter (BD). EGFP-Tat and EGFP were immunoprecipitated from U2OS cells using 25 μl of GFP-Trap magnetic beads (ChromoTek GmbH, Planegg-Martinsried, Germany) according to the manufacturer’s instructions.

Control of immunoprecipitation was carried out using immunoblotting. Beads were resuspended in the Laemmli sample buffer, boiled for 3 min, resolved on a 12.5% SDS-polyacrylamide gel and transferred to a nitrocellulose membrane. The membranes were blocked in 1% bovine serum albumin and incubated with either a monoclonal antibody against GFP (1:3000; Evrogen, Moscow, Russia) or a monoclonal antibody against B23 (1:10,000; Sigma-Aldrich, St. Louis, MO, USA) and monoclonal antibody against *β*-tubulin (1:10,000; Sigma-Aldrich, St. Louis, MO, USA). The membranes were washed three times with PBS (5 min each time) and were then incubated with secondary peroxidase-conjugated antibody (1:15,000; Sigma-Aldrich, St. Louis, MO, USA). The antibody-bound proteins were detected using Pierce ECL western blotting substrate (Thermo Scientific, Waltham, MA, USA), and images were acquired using a Gel DocXR system (Bio-Rad, Hercules, CA, USA).

For the pulldown assay, immobilized EGFP or EGFP-Tat was incubated with total RNA from U2OS cells for 2 h at 4°C on a rotary shaker. After three washing steps, the RNA was labeled with SYBR-Gold dye (Invitrogen). The beads were imaged using an AxioVert 200M microscope (Carl Zeiss, Oberkochen, Germany) equipped with an ORCAII-ERG2 cooled CCD camera (Hamamatsu Photonics K.K., Hamamatsu City, Shizuoka, Japan).

### NLS/NoLS bioinformatic analysis

We used 106 experimentally annotated NLSs of 88 viral proteins and 269 experimentally annotated NLSs of 228 human proteins (Tables S3). The dataset of the NLSs of human proteins was published elsewhere (11), and NLSs described in recent papers were included (Table S3). All NLSs of viral proteins were found via a search of UniprotKB database or published papers (Table S3). The NoLS dataset consisted of 24 experimentally determined NoLSs in 23 viral proteins and 71 NoLSs in 65 animal proteins (Table S4).

The presence of potential NoLSs in the human and viral protein datasets with experimentally determined NLSs was predicted by the NOD server (67), and the putative NLSs of the proteins in the NoLS dataset were predicted by NLStradamus (47).

The diversity and number of clusters of positive amino acids in an NLS were evaluated by custom R-script. The cluster was defined as a region in the NLS in which arginine residues and/or lysine residues/histidine residues were interspersed with another amino acid (there may be several such gaps, but each gap was not to be greater than one amino acid).

### Artwork

The contrast and brightness of the final images were adjusted using Adobe Photoshop (Adobe, San Jose, CA, USA). The gamma value was adjusted only for several panels, and these cases are noted in the figure legends. Figure 9 was created with BioRender.com.

## Code availability

R notebooks and custom scripts are available at https://github.com/lisitsynaom/NLS_Tat.

## ACKNOWLEDGEMENTS

We would like to thank S. Dokudovskaya, P.V. Lidsky, A.A. Zharikova and A.A. Mironov for stimulating discussions and valuable suggestions. A.O.Z. is grateful to Dr. G. Armeev for the comments on the quality assessment of crystal structures. We are grateful to Dr J. Borowiec for EGFP-NCL plasmid. The following reagents were obtained through the NIH AIDS Reagent Program, Division of AIDS, NIAID, NIH: HIV-1 HXB2 GST-Tat Expression Vector (GST-Tat 1 (86R)) from Dr. Andrew Rice (cat# 2367) and HIV-1 LTR lacZ Reporter Vector (pHIVlacZ) from Dr. Joseph Maio (cat #151). Docking experiments were carried out using the equipment of the shared research facilities of HPC computing resources at Lomonosov Moscow State University. The microscopy facility became available in the framework of the Moscow State University Development Program PNR 5.13. This work was supported by the Russian Science Foundation (grant 21-74-20134 to E.V.S.) and the Russian Foundation for Basic Research (grant 18-29-08012 to A.O.Z.).

E.V.S. conceived and designed the study. M.A.K., E.A.A., A.V.T., M.Y.S., M.A.T., D.M.P., A.I.K., Y.R.M. and E.V.S. performed cloning and microscopy studies. A.O.Z. and A.V.G. performed molecular docking. G.B. performed refinement of the crystal structure. O.M.L. and E.V.S. performed bioinformatic analysis. M.A.K., A.O.Z., O.M.L., A.V.G., Y.S.V. and E.V.S. analysed the data and wrote the manuscript.

